# Multi-omics analyses reveal novel effects of PLCγ2 deficiency in the mouse brain

**DOI:** 10.1101/2023.12.06.570499

**Authors:** Sarah C. Hopp, Juliet Garcia Rogers, Sabrina Smith, Gabriela Campos, Henry Miller, Savannah Barannikov, Eduardo Gutierrez Kuri, Hu Wang, Xianlin Han, Kevin F. Bieniek, Susan T. Weintraub, Juan Pablo Palavicini

**Affiliations:** Glenn Biggs Institute for Alzheimer’s and Neurodegenerative Diseases, University of Texas Health Science Center San Antonio; Department of Pharmacology, University of Texas Health Science Center San Antonio; Sam and Ann Barshop Institute for Longevity and Aging Studies, University of Texas Health Science Center San Antonio; Costa Rica Institute of Technology (TEC); Department of Pathology and Laboratory Science, University of Texas Health Science Center San Antonio; Department of Biochemistry & Structural Biology, University of Texas Health Science Center San Antonio; Department of Medicine, University of Texas Health Science Center San Antonio

**Author notes:** Corresponding Authors: Sarah C. Hopp and Juan Pablo Palavicini, 7703 Floyd Curl Drive, San Antonio, Texas, USA **Email:** and. **Author Contributions:** Sarah C. Hopp (Conceptualization, Methodology, Investigation, Formal Analysis, Visualization, Supervision, Project administration, Funding Acquisition, Writing – Original Draft, Writing – Review & Editing), Juliet Garcia Rogers (Investigation, Formal Analysis, Writing – Review & Editing), Sabrina Smith (Investigation), Gabriela Campos (Investigation), Henry Miller (Software, Formal Analysis, Visualization, Writing – Original Draft), Savannah Barannikov (Resources, Investigation), Eduardo Gutierrez Kuri (Investigation, Formal Analysis, Writing – Review & Editing), Hu Wang (Investigation), Xianlin Han (Resources, Supervision), Kevin F. Bieniek (Resources, Supervision), Juan Pablo Palavicini (Conceptualization, Methodology, Investigation, Formal Analysis, Visualization, Supervision, Project administration, Funding Acquisition, Writing – Original Draft, Writing – Review & Editing). **Competing Interest Statement:** The authors have no competing interests to disclose.

**Keywords:** Alzheimer’s disease, PLCG2, microglia, lipids, myelin

## Abstract

Phospholipase C gamma-2 (PLCγ2) catalyzes the hydrolysis of the membrane phosphatidylinositol-4,5-bisphosphate (PIP_2_) to form diacylglycerol (DAG) and inositol trisphosphate (IP_3_), which subsequently feed into numerous downstream signaling pathways. PLCG2 polymorphisms are associated with both reduced and increased risk of Alzheimer’s disease (AD) and with longevity. In the brain, PLCG2 is highly expressed in microglia, where it is proposed to regulate phagocytosis, secretion of cytokines/chemokines, cell survival and proliferation. We analyzed the brains of three-month-old PLCγ2 knockout (KO), heterozygous (HET), and wild-type (WT) mice using multiomics approaches, including shotgun lipidomics, proteomics, and gene expression profiling, and immunofluorescence. Lipidomic analyses revealed sex-specific losses of total cerebrum PIP_2_ and decreasing trends of DAG content in KOs. In addition, PLCγ2 depletion led to significant losses of myelin-specific lipids and decreasing trends of myelin-enriched lipids. Consistent with our lipidomics results, RNA profiling revealed sex-specific changes in the expression levels of several myelin-related genes. Further, consistent with the available literature, gene expression profiling revealed subtle changes on microglia phenotype in mature adult KOs under baseline conditions, suggestive of reduced microglia reactivity. Immunohistochemistry confirmed subtle differences in density of microglia and oligodendrocytes in KOs. Exploratory proteomic pathway analyses revealed changes in KO and HET females compared to WTs, with over-abundant proteins pointing to mTOR signaling, and under-abundant proteins to oligodendrocytes. Overall, our data indicate that loss of PLCγ2 has subtle effects on brain homeostasis that may underlie enhanced vulnerability to AD pathology and aging via novel mechanisms in addition to regulation of microglia function.

**Significance Statement:** The *PLCG2* gene contains a number of rare variants linked with increased and decreased risk for Alzheimer’s disease and longevity, but little is known about the role of PLCγ2 in normal brain function. The results described herein are significant because they describe the effects of knockout of PLCγ2 on brain cell types, thus mimicking the loss of function Alzheimer’s disease risk mutation. Our data describe novel effects of PLCγ2 deficiency on myelin homeostasis and mTOR signaling that have not been previously described that may underlie its association with Alzheimer’s disease pathogenesis and longevity.

## Introduction

Multiple *PLCG2* gene polymorphisms have been recently associated with Alzheimer’s disease (AD), including rare coding variants that increase (e.g. M28L, rs61749044) and reduce (e.g. P522R, rs72824905) risk of Alzheimer’s disease (AD) and increase the likelihood of longevity (P522R) (1–10). PLCG2 expression has also been associated with both amyloid and tau pathology (4, 11). The phospholipase C-γ-2 (PLCγ2) P522R protective variant is a functional hypermorph (12–16), while it is predicted that M28L is a loss-of-function variant (17). Expression of PLCγ2 M28L in the mouse brain led to a transcriptomic profile that correlated with human AD modules and led to increased microglia density when fed with a high fat diet (17). Other more strongly hypermorphic PLCG2 variants are associated with autoimmune PLCγ2-associated antibody deficiency and immune dysregulation (APLAID) (15), while other loss-of-function PLCG2 variants are associated with common variable immunodeficiency (CVID) and PLAID (18). However, the consequences of PLCγ2 loss-of-function in the brain remain largely unknown.

In the brain, PLCγ2 is primarily expressed by microglia, where it serves as an important signaling node downstream of TREM2 and CD33 (19, 20), both also AD risk genes (21–23). PLCγ2 catalyzes the hydrolysis phosphatidylinositol-4,5-bisphosphate (PIP_2_) to diacylglycerol (DAG) and inositol trisphosphate (IP_3_). DAG and IP_3_ feed into downstream signaling pathways involving protein kinase C (PKC), lipid biosynthesis, and intracellular calcium regulation. The TREM2/PLCγ2 signaling axis has been proposed to regulate phagocytosis, secretion of cytokines and chemokines, cell survival and proliferation (14, 24, 25). In the studies described herein, we tested the hypothesis that haploinsufficiency or full deficiency of PLCγ2 would lead to disrupted PIP_2_ and DAG metabolism in the mouse brain as measured by shotgun lipidomics and that these changes would result in changes in microglia phenotype as measured by gene expression profiling, proteomics and immunohistochemistry.

## Results

### PLCγ2 depleted mice display normal body and brain weight despite reduced survival, splenomegaly, and altered basal glucose homeostasis

Fourteen percent of male and six percent of female KO mice survived until weaning (Figure 1A). In other words, one out of two Plcg2 KO males survived until weaning age, while only one out of five Plcg2 KO females did so. Some Plcg2 KO mice died within the first several days postnatally. Average litter sizes at birth (5.2 pups) and at weaning (4.6 mice) indicate that ∼0.6 pups/litter died postnatally (Figure 1A). The number of animals that died post-natally prior to weaning, accounted for only about half the number of animals expected to have died based on Mendelian genetics, suggesting that about 50% of Plcg2 KO mice died in utero. These numbers are robust as they are based on 98 litters. Despite lower birth/weaning survival, all weaned Plcg2 KO mice survived until three months of age (end point of the study) and displayed normal body weights (Figure 1B). Brain dry weights were similar between genotypes (Figure 1C), which suggest that the premature death is not due to gross CNS abnormalities. On the other hand, spleen dry weight significantly increased in Plcg2 KO mice by 40% compared to WT controls (Figure 1D). Moreover, we found that Plcg2 KO mice display a trending reduction (p=0.082) by approximately −18% in non-fasted blood glucose compared to WT littermate controls as measured by 1-way ANOVA with Tukey post-hoc (Figure 1D); these trends reach significance when we include mice that were used as saline-treated controls for another study (Supplemental Figure 1D). Using such saline-treated controls enabled us to explore for potential sex differences. Similar trends were observed for both sexes (Supplemental Figure 1A-C). Note that Plcg2 HET mice displayed intermediate spleen weights and basal glucose levels for both sexes.

**Figure 1:**
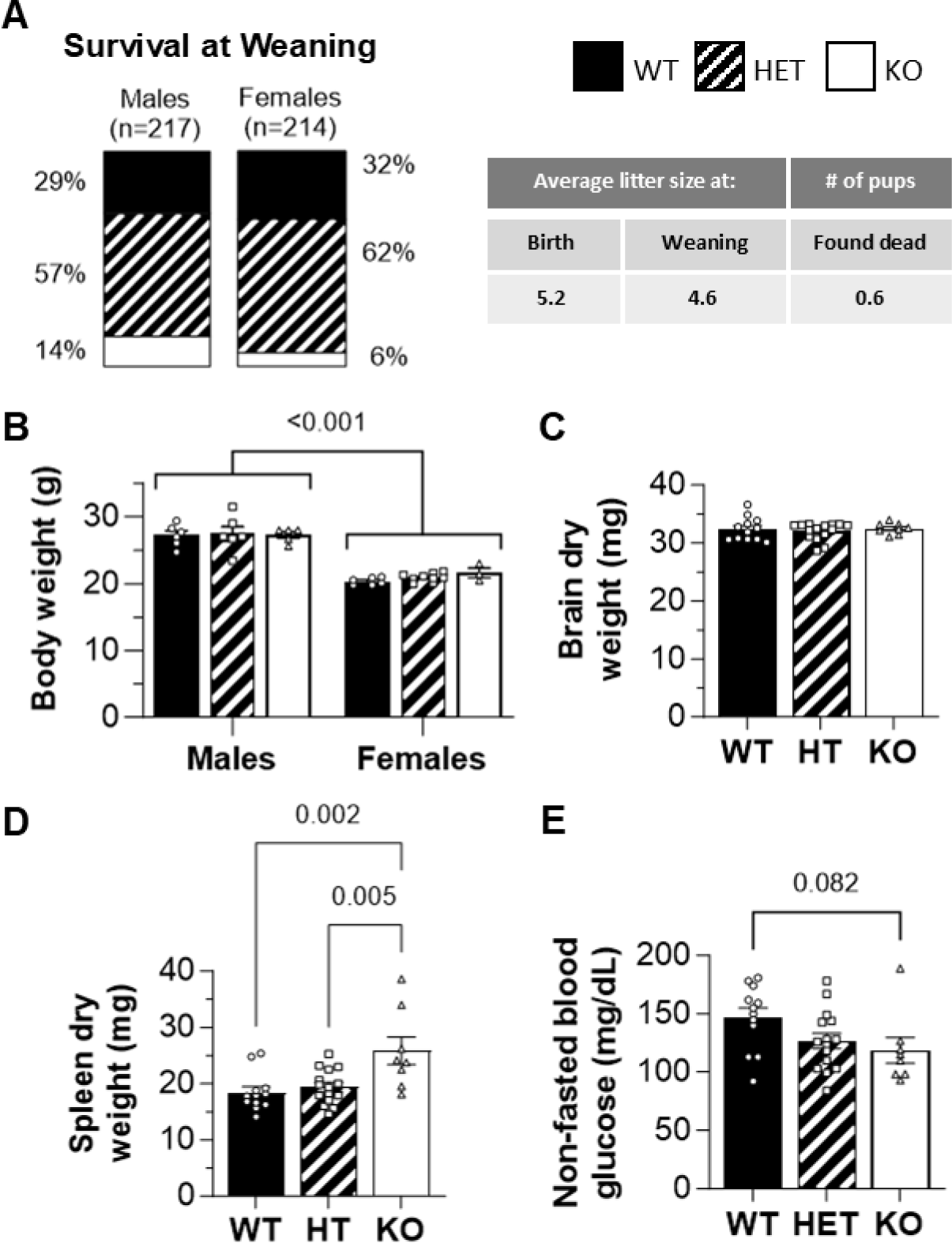
Basic phenotyping of PLCG2 WT, HET, and KO mice. A. Survival at weaning. 14% of male and 6% of female KO mice survived until weaning. PLCG2 KO females displayed higher mortality rates than PLCG2 KO males. B. Body weight. PLCG2 WT, HET, and KO mice display equivalent body weights with expected lower body weight in females. C. Brain weight. Brain dry weight was unchanged between genotypes. D. Non-fasting blood glucose. PLCG2 KO mice display a trending reduction in non-fasted blood glucose compared to WT mice (p=0.082); 2-way ANOVA (sex x genotype) revealed trending effects of sex (p=0.099) and genotype (p=0.066). *p<0.05 *** p<0.001

### PLCγ2 depletion alters the brain lipidome, with notable decreases in cerebroside and PIP_2_ species

Shotgun lipidomics on brain cerebrum revealed major sex-specific alterations and relatively minor genotype effects at the whole lipidome level (Figure S2A). Sex-specific principal component analyses (PCA) using 200+ lipid species revealed mild separation between Plcg2 KO and WT/HT mice in both sexes (Figure 2A-B). Similarly, unsupervised dendrogram analysis using Spearman distance and Ward algorithm generated two major clades, one composed exclusively of Plcg2 WT and HET mice, while a separate clade was composed primarily of KO mice (Figure S2B-C). A similar pattern was observed using supervised partial least squares-discriminant analysis (PLS-DA), where Plcg2 WT and KO mice were clearly clustered separately, while HET partially (females) or largely (males) overlapped with WT mice (Figure S2D-E). Volcano plots revealed 8 differentially abundant lipids (DALs) (Benjamini, Krieger and Yekutieli adjusted p-value <0.05,) between Plcg2 KO and WT (Figure 2C), 10 DALs between KO and HET (Figure 2D), and no DALs between HT and KO females (Figure S2F). Male results were similar, but the number of DALs was substantially smaller (Figure S2G-H), despite a larger n number. Most DALs were under-abundant in Plcg2 KO brains for either sex. Notably, most under-abundant DALs were myelin-specific or enriched lipid species, among them two cerebroside species (including CBS N24:1, the most abundant cerebroside species in the brain) were cosistently reduced in KO vs WT and/or HT mice in both sexes (Figure 2E). Similarly, exploratory unsupervised heatmaps displaying all metabolites with unadjusted p<0.1 (ANOVA comparing the 3 genotypes), clustered WT and HET mice together, while KO mice branched to a separate clade (Figure 2F-G).

**Figure 2:**
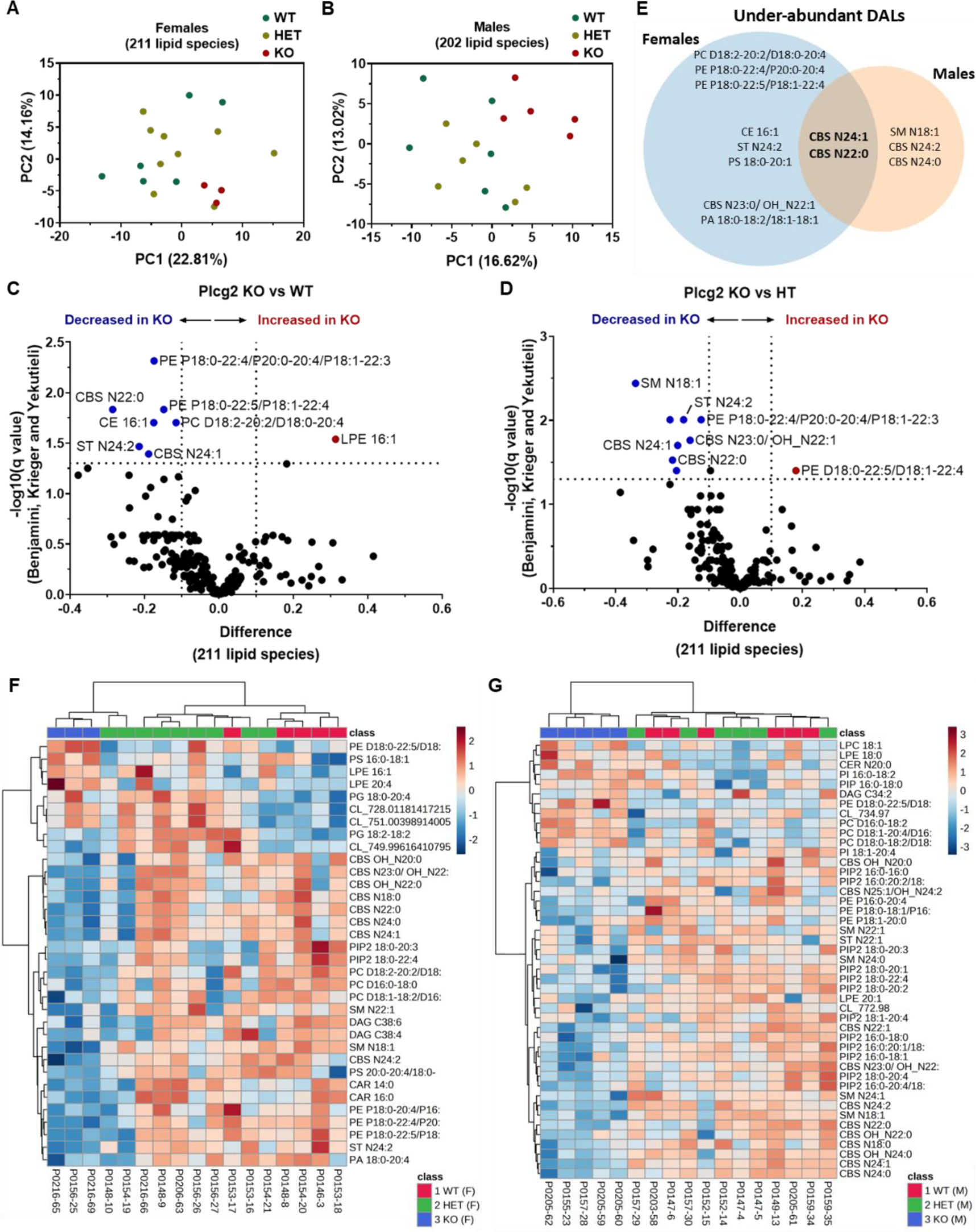
Lipidomic analysis. PCA analysis using cerebrum lipidomics data for female (A) and male (B) PLCG2 WT, HET, and KO mice. Volcano plot showing differentially expressed lipids between WT and KO (C) and between HET and KO (D) PLCG2 female mice. E. Ven diagram displaying the lipid species that were significantly reduced in PLCG2 KO compared to WT and/or HT mice in both sexes. F. Heat map showing top 50 differentially expressed lipids in WT, HET, and KO PLCG2 mice.

We next performed 2-way ANOVAs (genotype x sex) with post-hoc Tukey tests to examine changes in myelin lipid classes and subclasses, focusing on cerebrosides, sulfatides, and ethanolamine plasmalogens. We found a significant decrease of cerebrosides in the cerebrum that was dependent on genotype and sex, with post-hoc analysis revealing significantly less total cerebroside in Plcg2 KO mice compared to both WT and HET mice of both sexes (Figure 3A). Further characterization revealed significant effects of genotype and sex on decreasing N- and OH-cerebrosides levels in the cerebrum (Figure 3B-C), but only N-cerebrosides were significantly reduced in Plcg2 KO mice compared to WT and HET mice of both sexes (Figure 3B). There were no significant changes in total cerebrum sulfatide (Figure 3D), but further examination of N- and OH-sulfatide revealed a trending decrease in PLCG2 KO compared to WT mice (p=0.13, Figure 3E) and a main effect of sex (Figure 3F), respectively. There were significant effects of sex (p<0.001) and genotype (p=0.03) on total cerebrum ethanolamine plasmalogen, with a trending decrease in PLCG2 KO compared to WT mice (p=0.10, Figure 3G).

**Figure 3:**
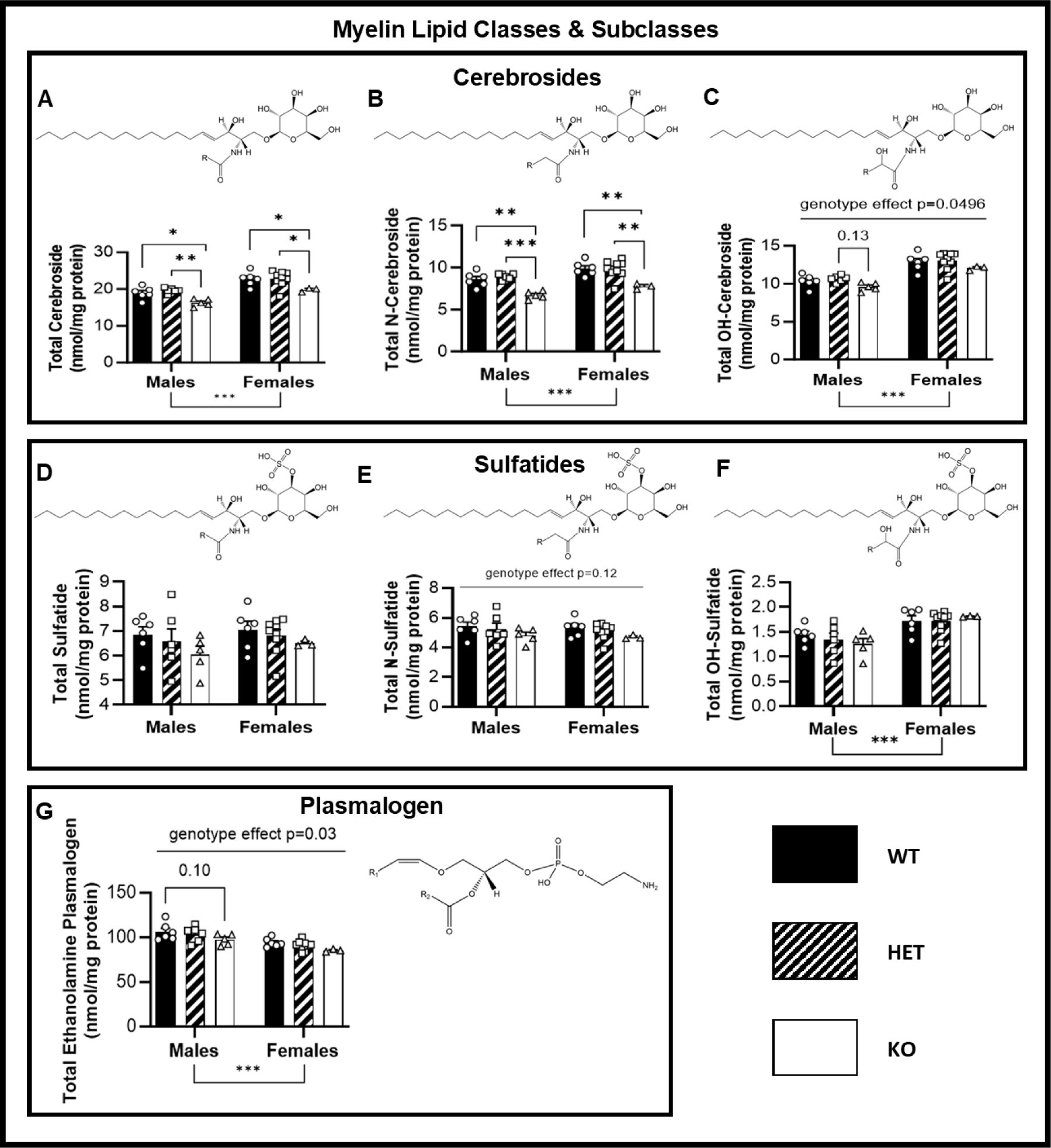
Myelin lipids. A. Total cerebrosides in the cerebrum were reduced in PLCG2 KO mice compared to both WT and HET mice of both sexes. B. N-cerebrosides levels in the cerebrum were significantly reduced in PLCG2 KO mice compared to WT and HET mice of both sexes. C. OH-cerebroside levels were significantly different between genotypes (2-way ANOVA genotype effect Tukey adj p=0.0496). Although multiple comparisons did not reach significance, PLCG2 KO mice tended to have slightly lower OH-cerebroside levels than WT and HET mice in both sexes. Note that cerebroside levels, including N- and OH-cerebrosides, were significantly higher in females compared to males across all genotypes. D. Total cerebrum sulfatide was unaltered by genotype or sex. E. N-sulfatide in the cerebrum trended to be altered by genotype (2-way ANOVA genotype effect p=0.12). Although multiple comparisons did not reach significance, PLCG2 KO mice tended to have slightly less N-sulfatide levels compared to WT and HET mice in both sexes. F. OH-sulfatide was not altered by genotype but was significantly higher in females (2-way ANOVA genotype effect p<0.001). G. Total cerebrum ethanolamine plasmalogen levels were significantly different between genotypes (2-way ANOVA genotype effect p=0.034). Although multiple comparisons did not reach significance, PLCG2 KO mice tended to have slightly lower ethanolamine plasmalogen levels than WT and HET mice in both sexes. Ethanolamine plasmalogen also varied by sex with females displaying higher levels than males.

Since the main function of PLCγ2 protein is to catalyze the hydrolysis of the membrane phospholipid PIP_2_ to form DAG and IP_3_, we closely examined changes in these lipid subclasses in Plcg2 WT, HET, and KO mice. We found a main effect of sex and a Plcg2 gene-dose dependent trending decrease in total DAG in the cerebrum (p=0.076, Figure 4A). We found similar sex differences in total saturated and monounsaturated DAG (Figure 4B-C) but no changes in total polyunsaturated DAG (Figure 4D). We also found no changes in total PIP or PIP_3_ levels in the cerebrum (Figure 4E and 4G), but we did observe a significant main effect of genotype, with a specific loss of PIP_2_ in Plcg2 KO males compared to sex-matched WT and HET mice and a decreasing trend in females (Figure 4F).

**Figure 4:**
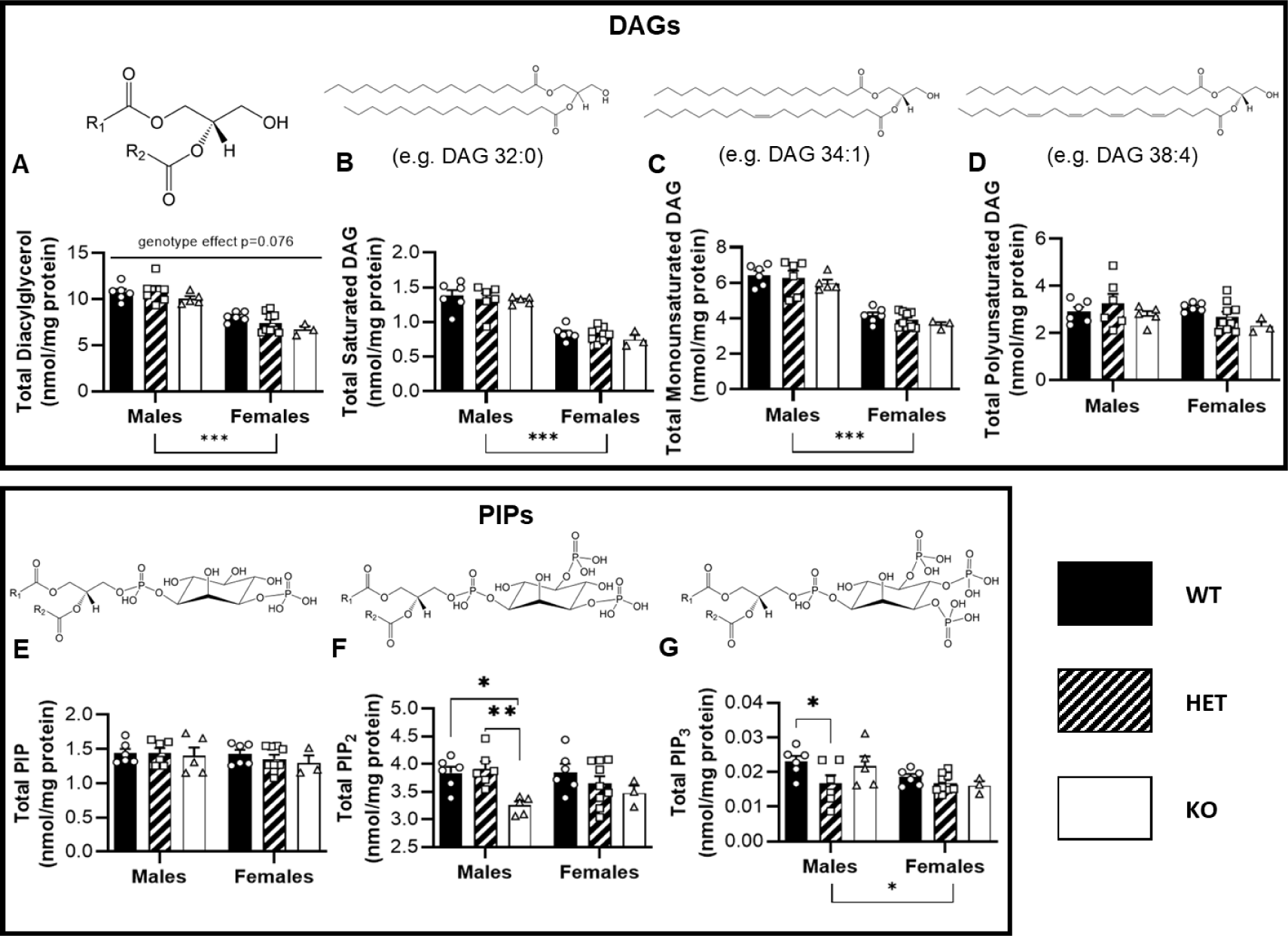
DAG and PIP. A. Total DAG in the cerebrum was reduced in females and trended towards reduction with reduced PLCG2 gene (p=0.076). B. Total saturated DAG was significantly reduced in female mice C. Total monounsaturated DAG was significantly reduced in female mice D. There were no changes in total polyunsaturated DAG. E. There were no changes in total PIP in the cerebrum. F. PIP2 was significantly reduced with reduced PLCG2 gene dose with a specific loss of PIP2 in PLCG2 KO males compared to sex-matched WT and HET mice. G. Total PIP3 levels in the cerebrum was significantly reduced in female mice and there was a trending (p=0.0326) change by genotype with male HET mice having significantly lower PIP3. *p<0.05 **p<0.01 *** p<0.001

### PLCγ2 depletion changes oligodendrocyte-, astrocyte- and microglia-related gene expression

We used the Nanostring nCounter^®^ Glial Profiling Panel to assess changes in gene expression of 770 mouse genes across 50+ pathways involved in glial cell biology in the cerebrum and brain stem of Plcg2 WT, HET and KO male and female mice. Principal component analysis (PCA) revealed, as expected, substantial sex- and brain region-specific effects on the expression of the Glial Profiling Panel before (Supplemental Figure 2A-B) and after (Figure 5A-B) quality control/noise reduction. Although unbiased dimensionality reduction of the metadata did not reveal genotype effects at the whole Glial Profiling Panel level, i.e. Plcg2 KO did not clearly separate from Plg2 WT or HET mice on the PCA plot (Supplemental Figure 2C and Figure 5C), pairwise differential gene expression analysis on the metadata (including both brain regions and both sexes) revealed 17 differentially expressed genes (DEGs) between Plcg2 WT and KO (Figure 5D, Supplemental Dataset 3) and 14 DEGs between Plcg2 HET and KO using *DESeq2* (Wald’s test, Benjamini Hochberg, p-adj<0.05) (Figure 5E, Supplemental Dataset 4). A large portion of the overexpressed DEGs between those two comparisons were shared (Figure 5F), while no underexpressed DEGs were shared (Supplemental Figure 2D). No DEGs were obtained between Plcg2 WT and HET mouse brains (Supplemental Figure 2E, Supplemental Dataset 5). Wald test statistics across genotype comparisons, revealed a strong agreement between “Plcg2 KO vs WT” and “Plcg2 KO vs HET” (Figure 5G), but not between other comparisons (Supplemental Figure 2F-G). Taken together our metadata analysis demonstrated that Nanostring RNA expression profiles in Plcg2 WT and HET brains were virtually identical, while a subset of genes was differentially expressed in Plcg2 KO compared to Plcg2 WT and HET.

**Figure 5:**
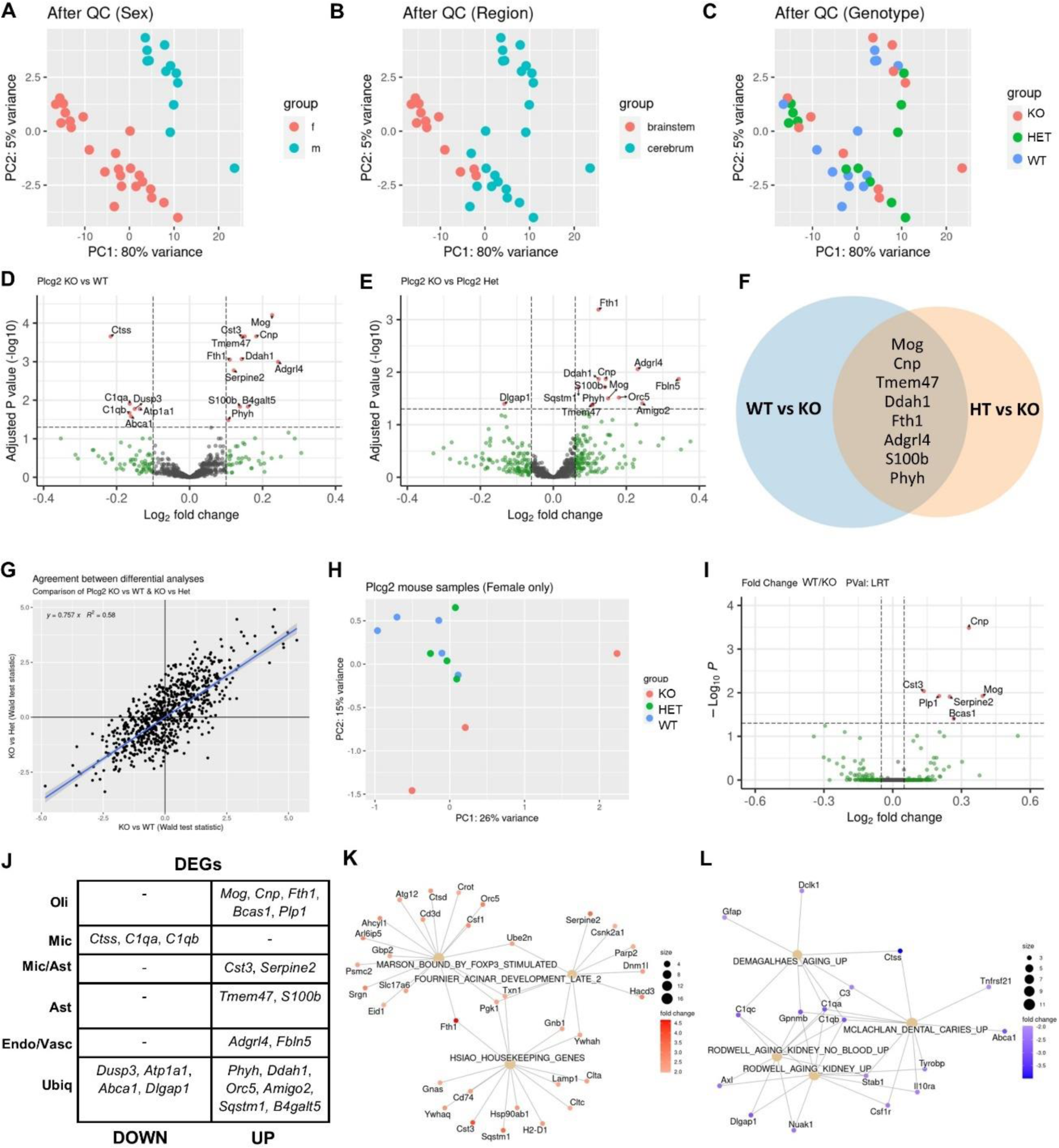
Nanostring glia profiling panel. A. PCA showing sex-specific effects on gene expression. B. PCA showing brain region-specific effects on gene expression. C. PCA showing limited effect of genotype on gene expression. D. Volcano plot illustrating differentially expressed genes between female WT and KO mice. E. Volcano plot illustrating differentially expressed genes between male WT and KO mice. F. Venn diagram showing shared DEGs between WT vs. KO and HET vs. KO. G. Correlation analysis of Wald test statistics show agreement between “Plcg2 KO vs WT” and “Plcg2 KO vs HET”. H. Sex-specific PCA plot for female cerebrum samples only show separation between Plcg2 KO and Plcg2 WT/HET gene expression. I. DESeq2 likelihood ratio test (LRT, Benjamini-Hochberg) differential gene expression analysis revealed six DEGs in female cerebrum. J. Upregulated and downregulated DEGs by cell type. K. Pathway analysis using Molecular Signatures Database (MSigDB) identified three significantly over-expressed pathways and K. four significantly under-expressed pathways (Figure 5L) using the Wald statistic.

To test for potential sex-specific effects between Plcg2 genotypes at the RNA profiling level, we performed exploratory subsequent analysis on the subset of samples that were balanced between both sexes, i.e., cerebrum samples, which were also used for lipidomic and proteomic analysis. Sex-specific PCA plots revealed clear separation between Plcg2 KO and Plcg2 WT/HET female brains (Figure 5H), but not for males where there was no clear separation between genotypes (Supplemental Figure 2H). Furthermore, DESeq2 likelihood ratio test (LRT) differential gene expression analysis revealed six female DEGs (Figure 5I, Supplemental Dataset 6) but not a single male DEG (Supplemental Figure 2I). Note that sex-specific analyses were underpowered and unadjusted.

Several DEGs that were under-expressed in Plcg2 KO brains on our metadata were microglia genes, including complement components (*C1qa* and *C1qb*) and a lysosomal proteinase that participates in the degradation of antigenic proteins to peptides for presentation on MHC class II molecules (*Ctss*) (Figure 5J). Genes coding for major myelin proteins (*Mog*, *Cnp*, *Plp1*) were also significantly altered in metadata and/or sex-specific analysis, as well as other genes primarily expressed by oligodendrocytes (*Fth1*, *Bcas1*) (Figure 5J). Other up-regulated DEGs are primarily or highly expressed by astrocytes (*Tmem47*, S100b, *Cst3*, and *Serpine2*). Considering that microglia depletion leads to compensatory upregulation of astrocytic genes and activity (26, 27), taken together these results strongly suggest that Plcg2 deficiency leads to a mild inactivation of microglia. Finally, upregulated DEGs also included *Adgrl4*, primarily expressed in endothelial cells; and *Fbln4*, primarily expressed in vascular fibroblast-like cells (Figure 5J). Exploratory pathway analysis using genes that were significantly altered (unadjusted p value<0.05) in at least two genotype comparisons and the Molecular Signatures Database (MSigDB) identified three significantly over-expressed pathways (Figure 5K) and four under-expressed pathways (Figure 5L) using the Wald statistic. The most relevant over-expressed pathway involves genes with promoters bound by FOXP3, a transcription factor involved in regulatory T-cell function, in stimulated hybridoma cells(28). On the other hand, three under-expressed pathways were related to aging, consistent with the previously reported association between PLCG2 and longevity(2).

Moreover, we used the Nanostring advanced analysis tool to examine different pathways altered by Plcg2 gene dose and sex. We found a significant main effect of genotype on genes related to homeostatic microglia (p=0.001, Supplemental Figure 3A), which were significantly reduced in female KO brains compared to those of WT/HT, and trended to decrease in male KOs. Similarly, we found decreasing trends in markers of neurodegenerative microglia (MgND, p=0.081, Supplemental Figure 3B) and stage 1 disease associated microglia (DAM) markers (p=0.178, Supplemental Figure 3C). We also found interaction between sex and genotype in MAPK and PI3K related genes (p=0.028, Supplemental Figure 3D), T-cell signaling (p=0.046, Supplemental Figure 3E), and complement system genes (p=0.046, Supplemental Figure 3F). There were also sex-specific effects on expression of oligodendrocyte (p<0.001, Supplemental Figure 3G), macrophages/microglia (p<0.001, Supplemental Figure 3H), and neuron (p<0.001, Supplemental Figure 3I) genes.

### PLCγ2 depletion reduces microglia and myelin density but does not alter neurons or astrocytes

Due to the observed changes in microglia gene expression, the area coverage and number of brain myeloid cells were evaluated by immunofluorescence (IF) for the pan-myeloid marker Iba1 (Figure 6A-C). There was no significant difference in the overall area coverage of Iba1 per section as quantified by automated thresholding (p=0.22, Figure 6D). However, automatic cell counting revealed a significant decrease in the density of cells per unit area in KO mice compared to WT mice (p<0.05, Figure 6E). Due to the observed changes in myelin lipids, the area fraction of MAG was evaluated by IF as a marker of myelin changes (Figure 7A-C). There was a trending main effect of genotype on MAG area fraction (p=0.14) with KO mice showing less MAG labeling compared to WT (p=0.16) and HET (p=0.24) (Figure 7D). We also evaluated changes in NeuN and did not observe any changes in the area fraction of neuronal nuclei across any genotype (Figure 7E) nor did we observe any gross changes in NeuN distribution or morphology. Finally, we evaluated the area fraction of astrocytes labeled with GFAP to see if increase in astrocytes could be compensating for the mild reduction in microglia (Figure 7F-H), but we found no effect of genotype on astrocyte area fraction (Figure 7I).

**Figure 6:**
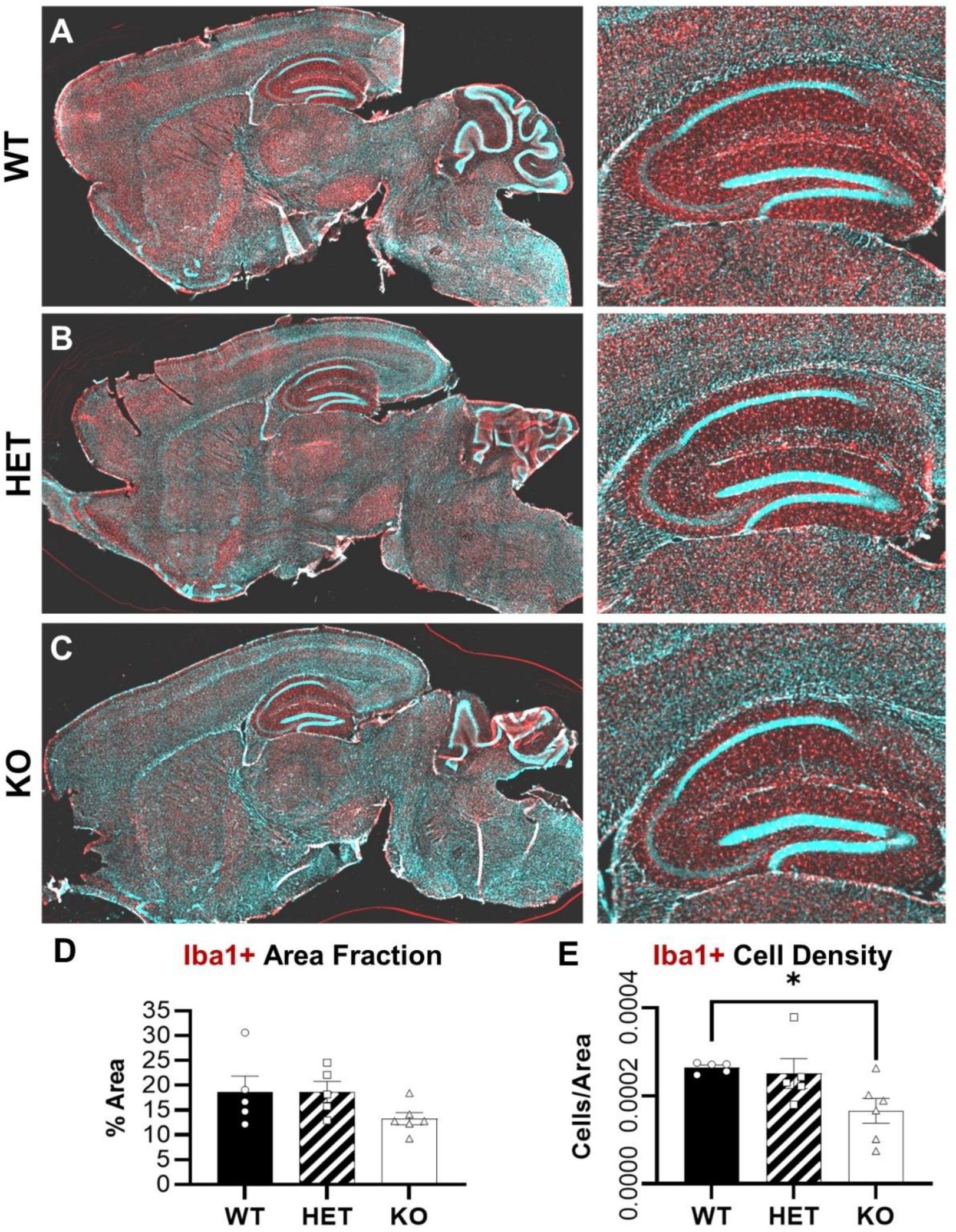
Iba1 immunohistochemistry. A-C. Microglia were visualized in sagittal sections with the pan-myeloid marker Iba1 (red) and counterstained with DAPI to label nuclei in A) WT, B) HET, and C) KO PLCG2 mice. Inset shows hippocampus. D. There was weak trend (p=0.22) towards reduced overall area fraction of Iba1+ staining in PLCG2 KO mice. E. The density of Iba1+ microglia was significantly reduced in PLCG2 KO mice. *p<0.05

**Figure 7:**
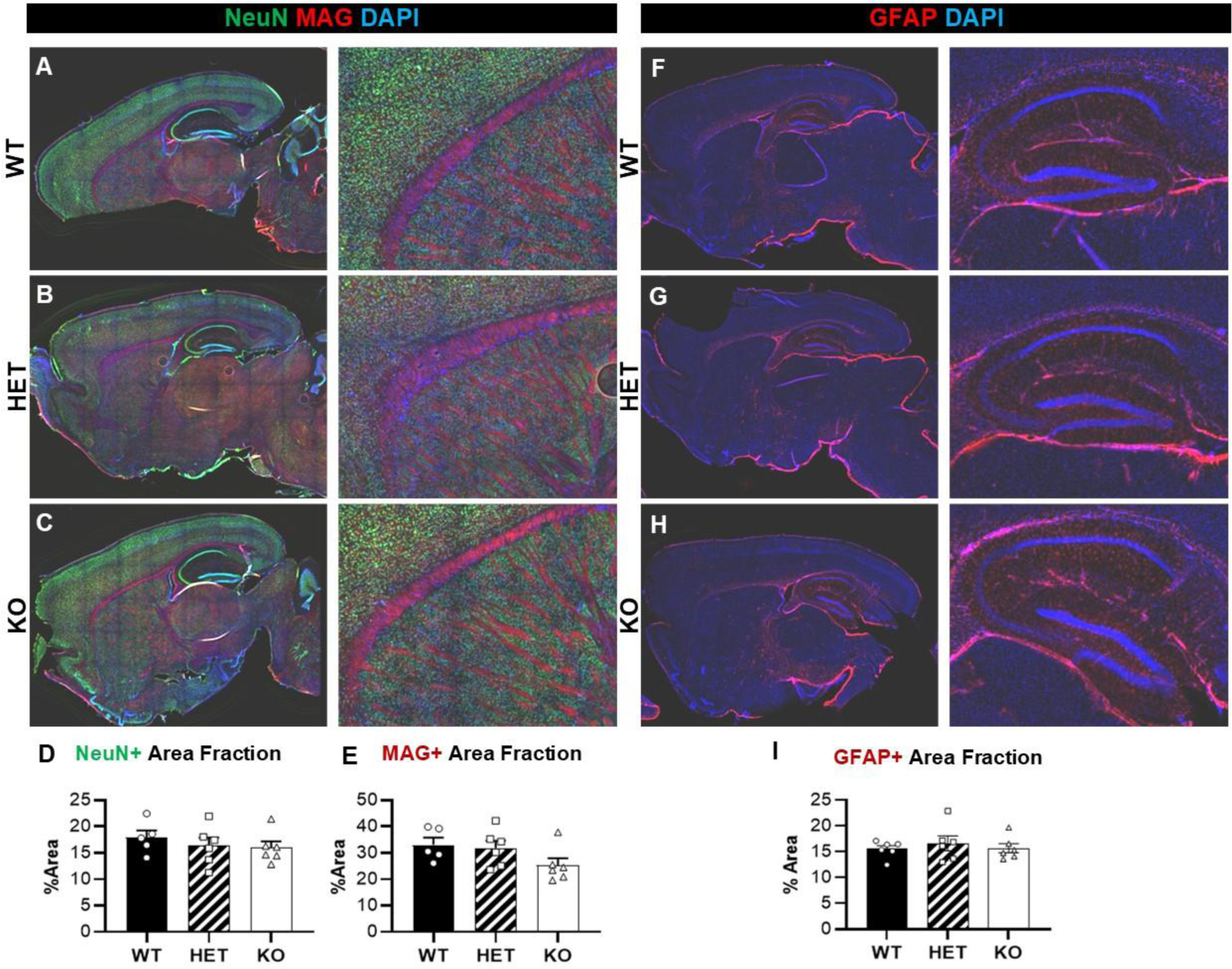
NeuN, MAG, and GFAP immunohistochemistry. A-C. White matter and neuronal nuclei were visualized in sagittal sections with MAG (red) and NeuN (green), respectively; nuclei were counterstained with DAPI in A) WT, B) HET, and C) KO PLCG2 mice. Inset shows white matter tracts. D. There was no difference in NeuN area fraction. E. There was a trending main effect of genotype on MAG area fraction (p=0.14) with KO mice showing less MAG labeling compared to WT (p=0.16) and HET (p=0.24). F-H. Astrocytes were visualized with GFAP (red) and counterstained with DAPI to label nuclei in F) WT, G) HET, and C) KO PLCG2 mice. I. There was no difference in GFAP area faction.

### Exploratory proteomics reveal altered amino acid metabolism in the brains of PLCγ2 depleted mice

We used unbiased proteomics to assess differentially abundant proteins (DAPs) in brain tissue from female Plcg2 KO, HET, and WT mice. Female mice were used here for proteomics analyses since most previous analyses indicated stronger effects in females. PCA revealed differences in proteins due to mouse and/or sample effects outweigh differences due to genotype (Figure 8A), indicating that Plcg2 depletion does not have major effects at the whole brain proteome level under physiological conditions in young adult mice. Correlating T-test statistics revealed weak correlations across differential analysis comparisons (Supplemental Figure 4A-C). Exploratory pairwise comparisons using unadjusted p values between the different genotypes revealed 552 DAPs (Figure 8D-F, Supplemental Datasets 7-9), including 356 over-abundant and 196 under-abundant proteins under Plcg2 deficient conditions. Notably, 102 DAPs were shared in two different genotype comparisons, 77 of which were increased and 25 decreased (Figure 8B-C). It is important to note that none of the 4215 proteins that were reliably detected and passed QC reached significance after applying Benjamini-Hochberg correction, so data is reported as unadjusted p-values for data exploration. To better understand how samples related to one another with respect to differential protein abundance, we performed hierarchical clustering using all DAPs, and found that each of the three genotypes clustered separate from each other (Figure 8G). Surprisingly, contrary to our lipidomics and gene profiling results, the brain proteome of HET mice was more similar to KO than WT. Furthermore, we performed KEGG pathway analysis using all DAPs to understand the biological processes that were perturbed across Plcg2 genotypes. The results highlighted a variety of pathways relevant to aging and neurodegenerative disease. Interestingly, the results also show evidence of alterations in amino acid metabolism and ribosome biogenesis (Fig. 8H). Similarly, mTOR signaling-related pathways were the only ones to reach significance (q-value<0.035) after performing BioCarta pathway analysis using overabundant proteins (Figure 8I). On the other hand, Enrichr Cell type analyses using underabundant proteins revealed oligodendrocytes and oligodendrocyte precursor cells, as the only cell types significantly altered (q-value<0.05) in Plcg2 KO brains (Supplemental Figure 4D-F).

**Figure 8:**
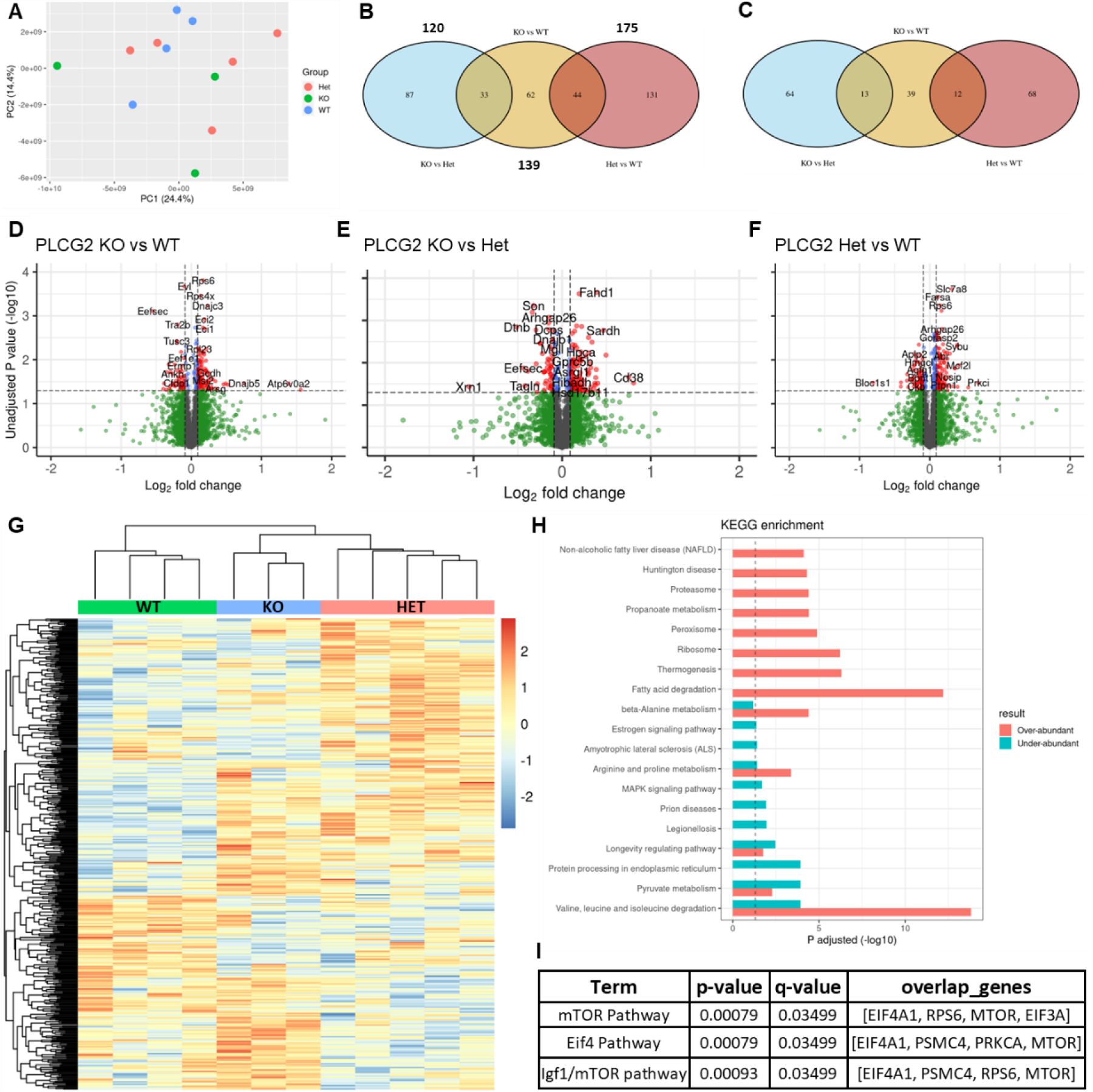
Proteomics analysis. A. PCA revealed differences in proteins due to mouse and/or sample effects largely outweigh differences due to genotype (Figure 8A). B-D. Correlating T-test statistics across genotype comparisons. B. “KO vs HET” and “KO vs WT” DAPs C. “Plcg2 KO vs WT” and “Plcg2 HET vs WT” DAPs D. “HET vs WT” and “KO vs HET” DAPs. E-G. Pairwise comparisons of DAPs between the different genotypes. E. DAPs between KO and WT. F. DAPs between KO and HET. G. DAPs between HET and WT. H. Hierarchical clustering shows that KO and HET samples clustered together, excluding WT. I. Venn diagram showing overall between pairwise DAPs by genotype. H. KEGG pathway analysis of over- and under-abundant DAPs. I. BioCarta pathway analysis using overabundant DAPs.

## Discussion

Yields of Plcg2 KO mice were lower than expected based on theoretical Mendelian genetics (∼25%) with females showing higher mortality rates than males. Taken together, our results imply that mortality is almost certainly driven by developmental defects. The cause of death was not determined but may be related to the previously described essential role of PLCγ2 in initiating the separation of the blood and lymphatic vasculature (29). Previous studies have reported internal hemorrhaging during development in other Plcg2 deficient mouse models (29, 30), which have been proposed to be caused by rupture of blood-filled fragile lymphatic vessels by mechanical stress or pressure of the blood flow (29). Although internal bleeding likely explains early mortality, the mechanism(s) underlying the observed sex-specific mortality rates remain unclear. Consistently, while performing gross organ examination during harvest, we noticed that the intestines of Plcg2 KO mice were clearly different, particularly in regards to their apparent vascular system, differences were noticeable before and after saline cardiac perfusion and were more evident in females. Specifically, we observed an evident increase in the density of blood-filled vessels in Plcg2 KO intestines, that were thinner and more arborized than those of WT controls. Additionally, Plcg2 deficiency led to a substantial enlargement of the spleen measured by dry weight. These results are in line with high levels of Plcg2 gene expression in spleen B-cells (particularly in T1, T2 and follicular B-cells) (Gene Expression Commons), suggestive a potential role of Plcg2 in spleen homeostasis. In fact, a previous study reported marked decreases in the most mature B cell population in Plcg2 deficient mice(30).Taken together, these data may suggest strong compensatory mechanisms on surviving Plcg2 deficient animals that could bias downstream analyses for highlighting compensatory mechanisms for survival rather than direct effects of PLCγ2 loss.

The main function of PLCγ2 protein is to catalyze the hydrolysis of PIP_2_ to produce DAG and IP_3_. Although we found expected decreases in one of the products of the PLCγ2 enzymatic reaction, i.e. DAG, in Plcg2 KO mice (Figure 3), we found a decrease in the substrate of PLCγ2 enzymatic reaction, i.e. PIP_2_ (Figure 3), rather than the expected accumulation. Our results are consistent with the ‘buffering’ of global plasma membrane PIP_2_ pool hypothesis(31), which proposes that stimulation of PIP_2_ hydrolysis (e.g. via PLCγ2) occurs concurrently with an activation of PIP_2_ synthesis via Ca^2+^-sensitive stimulation of PI4-kinase(32). Thus, loss of PLCγ2 would not only prevent PIP_2_ hydrolysis, but also prevent its synthesis, potentially explaining the observed overall reduction of PIP_2_ levels. Considering that the PLC family is composed by 14 members (including 2 catalytically inactive members), these results suggest that PLCγ2 is a major regulator of brain PIP_2_ homeostasis despite being expressed at relatively low levels in the brain compared to other family members. Our proteomics results identified 8 PLC family members in the mouse brain, with PLCβ1 being by far the most abundant one, and PLCγ2 being below detection limit. We could not find evidence of Plcg2 deficiency leading to altered levels of other PLC family members in our RNA profiling and proteomics studies. A previous study examining microglia gene expression from Plcg2 KO mice also found rewiring of PIP_2_ and DAG metabolism, with decreases in DGKE and DAGL which both degrade DAG and increases in PLA2G4A which shunts PI away from production of PIP_2_ to generate free fatty acids such as arachidonic acid (15). While we did not measure PLA2G4A, DGKE, or DAGL in our study, compensatory reductions in expression of these enzymes could explain our observed increase in PIP_2_. In alignment with our data, this study also found reduced production of DAG in microglia isolated from Plcg2 KO mice (15). Given the complex interconnectivity between lipid pathways, we think that Plcg2 depletion leads to an alteration in alternative pathways that produce or generate PIP_2_, explaining the observed results.

In microglia PLCγ2 activity has been documented to be downstream of TREM2 (19) and Fc receptors (15), regulating inflammatory processes, migration, and endocytosis (13, 16, 25, 33, 34). Our gene expression data partially align with previously published transcriptome data from neonatal microglia isolated from Plcg2 KO mice, which found reduced expression of disease-associated microglia markers, as well as reduced expression of genes associated with lysosomal function, phagocytosis of apoptotic cells, and extracellular matrix degradation and increased expression of complement, major histocompatibility complex, inflammatory chemokine, and interferon response genes (15). Consistent with the expected role of PLCγ2 in microglia and with a recent report that assessed gene expression profiles in a Plcg2-inactivated mouse model (11), all DEGs that were under-expressed in Plcg2 KO brains on our data were microglia genes (Figure 5J). Other up-regulated DEGs are primarily or highly expressed by astrocytes (*Tmem47*, S100b, *Cst3*, and *Serpine2*). Considering that microglia depletion leads to compensatory upregulation of astrocytic genes and activity (26, 27), taken together these results strongly suggest that Plcg2 deficiency leads to a mild inactivation of microglia. Differences between our results and these previously published results may be related to the age of the mice studied and methodological differences, e.g. isolated microglia vs whole brain gene expression.

Surprisingly, our lipidomics results also showed significant decreases in myelin-specific lipids (i.e. cerebrosides) and decreasing trends in myelin-enriched lipids (i.e. sulfatides and phosphatidylethanolamine plasmalogens) in Plcg2 KO mice (Figure 3). This reduction in myelin lipids corresponds with our observed trending reduction of MAG by immunohistochemistry, which could result from altered oligodendrocyte homeostasis and/or myelin abnormalities. Notably, oligodendrocytes and oligodendrocyte precursor cells came as top hits in exploratory Enrichr cell type analysis using under-abundant proteins. Consistent with our lipidomics and proteomics results, gene expression of major myelin proteins (*Mog*, *Cnp*, *Plp1*) and other genes primarily expressed by oligodendrocytes (*Fth1*, *Bcas1*) were also significantly altered in our Nanostring analysis. Counterintuitively, myelin/oligodendrocyte genes were actually increased, suggesting potential compensatory effects at the RNA level. These results were surprising given that microglia are the main cell type that express Plcg2 in the brain, with relatively low expression in oligodendrocytes, the cells that produce myelin (12). Microglia-oligodendrocyte interactions may account for this change in myelin lipids since microglia are critical for the development of myelin (35) and in the study herein Plcg2 is disrupted constitutively. Alternatively, there is some evidence that a subset of oligodendrocytes, termed immune oligodendrocytes, express PLCG2 mRNA (36–38), suggesting that perhaps the overall decreases in myelin lipids in Plcg2 KO mice may be due to cell autonomous effects of PLCG2 loss. Follow-up studies aimed at dissecting between these two possibilities are warranted. Notably, previous studies have shown that the lipids that were or tended to be decreased in PLCG2 KO mice (cerebrosides, sulfatides and plasmalogens) have been also reported to be decreased early in AD (39–41).

Taken together with our lipidomic, gene expression, and histology data suggest that loss of PLCγ2 alters microglia reactivity, potentially leading to downstream loss of myelin without strongly affecting neurons by the measures we have examined. PLCγ2 mRNA has relatively low expression in oligodendrocytes compared to microglia (12), so it is likely that microglia-oligodendrocyte interactions may account for this change in myelin lipids since microglia are critical for the development of myelin (35). However, other cellular interactions may mediate the disruption of myelin lipids in Plcg2 KO mice. PLCγ2 is necessary for separation of blood and lymphatic vessels in mice (29) and osteoclastogenesis (42), and PLCγ2 outside of the brain has roles in immune deficiencies, autoimmunity, and cancer (18). PLCγ2 is an important downstream mediator of B cell an T cell receptors (18). In the studies herein, we found a significant change in T cell signaling as measured by Nanostring gene expression analysis (p=0.046) with specific increases in male PLCγ2 KO mice (Supplemental Figure 3D). Moreover, exploratory pathway analysis using overexpressed genes in Plcg2 KO mice revealed a “FOXP3 stimulated” molecular signature (Figure 5K), implying a potential activation of regulatory T cells under PLCγ2 deficient conditions. Recent studies have shown that T cells mediate myelin dysfunction in aging via interferon signaling (43); perhaps loss of PLCγ2 enhances T cell mediated oligodendrocyte and microglia dyshomeostasis which could explain our results showing that myelin lipids decrease in PLCγ2 KO mice (Figure 3A-C). We observed increases in complement and decreases in homeostatic microglia in female Plcg2 KO mice (Supplemental Figure 3A, 3F). Interestingly, changes in microglia and T cells were in females and males respectively, suggesting sexually dimorphic effects of Plcg2 KO on different cell types. Sexual dimorphism was pronounced in survival of Plcg2 KO mice (Figure 1), as well as our lipidomic analyses, with changes more pronounced in females (Figure 2). The mechanisms of the sexually dimorphic effects of Plcg2 KO are unclear but understanding them could be important to understanding human health and disease; PLCγ2 plays a role in both AD and autoimmunity, both of which are more prevalent in women. Notably, our results also show changes in other genes associated with AD (*Bcas1*, *Abca1*) and pathways associated with longevity (mTOR signaling).

Overall, we found that depletion of PLCγ2 lead to subtle changes in microglia phenotype in the absence of inflammatory stimuli, suggestive of reduced microglial reactivity. Our data suggest a potential role for PLCγ2 in myelin, amino acid, and glucose homeostasis that requires further investigation. Overall, our data suggest that loss of PLCγ2 function has subtle but highly sexually dimorphic effects on brain homeostasis that may underlie enhanced vulnerability to AD pathology via microglia and, surprisingly, myelin dysfunction.

## Materials and Methods

### Mice

All procedures were approved by the Institutional Animal Care and Use Committee at University of Texas Health Science Center San Antonio. Experiments were performed on 3-month-old mice heterozygous and homozygous for PLCG2 KO (JAX #029910) and WT littermates of both sexes on a congenic C57BL6/J background (MMRRC #34848-JAX). Up to five mice were housed per cage with a 14:10 light: dark cycle, with lights on at 7am and off at 9pm. At weaning, mice were genotyped according to the protocol provided by Jackson Labs and ear notched for identification.

### Euthanasia and Tissue Processing

At 3 months of age, body weights were recorded and non-fasted blood glucose levels were measured using Contour glucose meter/strips (Bayer), mice were subsequently euthanized using isoflurane and perfused with PBS in accordance with American Veterinary Medical Association guidelines. Brains were hemisected and one hemibrain was post-fixed in 4% PFA and sliced into 40 µm sections using a Leica VT1000 S vibratome. The cerebrum and brainstem from the other hemibrain was snap-frozen in liquid nitrogen. Brain tissues were lyophilized for 48 hours at 0.5 mbar using a Labconco freeze dryer and weighted on an analytical balance.

### Shotgun Lipidomics

Cerebrum samples were homogenized in 0.1 PBS using Precellys^®^ ceramic beads/homogenizer equipped with a Cryolys Evolution cooling system (Bertin Technologies). Total protein concentrations were determined using bicinchoninic acid (BCA) protein assay (Thermo Fisher Scientific). Lipids were extracted by a modified procedure of Bligh and Dyer extraction in the presence of internal standards, which were added based on protein content as previously described(44). Lipids were assessed using a triple-quadrupole mass spectrometer (Thermo Scientific TSQ Altis) and a Quadrupole-Orbitrap™ mass spectrometer (Thermo Q Exactive™) equipped with a Nanomate device (Advion Bioscience Ltd., NY, USA) as previously described(45, 46). All full and tandem MS mass spectra were automatically acquired using a customized sequence subroutine operated under Xcalibur software. Data processing including ion peak selection, baseline correction, data transfer, peak intensity comparison, ^13^C deisotoping, and quantitation were conducted using a custom programmed Microsoft Excel macro after considering the principles of lipidomics(47). Data was analyzed using the MetaboAnalyst metadata module with genotype and sex as categorical variables. Data was log transformed and pareto scaled, top 25 lipid species by Two-way ANOVA (Type I) with post-hoc Bonferroni arranged by sex (first) and genotype (second). Volcano plots were generated with GraphPad Prism using multiple t-tests with post-hoc FDR 5% (2-stage step-up method, Benjamini, Kieger Yekutieli).

### Proteomics

Approximately 3 mg of each lyophilized mouse cerebrum sample were placed in a MicroTube fitted with a MicroPestle (Pressure BioSciences), and 30 µl of homogenization buffer [10% SDS in 50 mM triethylammonium bicarbonate (TEAB, Thermo Fisher Scientific) containing protease/phosphatase inhibitors (Halt, Thermo Fisher Scientific) and nuclease (Universal Nuclease, Pierce/Thermo Fisher Scientific). Tissue samples were homogenized in a Barocycler (Pressure BioSciences, Inc.) for 60 cycles at 35 °C and then the MicroTubes were centrifuged at 21,000 x g for 10 minutes. Aliquots corresponding to 100 µg protein (EZQ™ Protein Quantitation Kit; Thermo Fisher Scientific) were reduced with tris(2-carboxyethyl)phosphine hydrochloride (TCEP), alkylated in the dark with iodoacetamide and applied to S-Traps (mini; Protifi) for tryptic digestion (sequencing grade; Promega) in 50 mM TEAB. Peptides were eluted from the S-Traps with 0.2% formic acid in 50% aqueous acetonitrile and quantified using Pierce™ Quantitative Fluorometric Peptide Assay (Thermo Fisher Scientific).

Data-independent acquisition mass spectrometry was conducted on an Orbitrap Fusion Lumos mass spectrometer (Thermo Fisher Scientific). On-line HPLC separation was accomplished with an RSLC NANO HPLC system (Thermo Fisher Scientific/Dyonex). A pool was made of all of the samples, and 2-µg peptide aliquots were analyzed using gas-phase fractionation with staggered 4-m/z windows (30k resolution for precursor and product ion scans, all in the orbitrap). The 4-mz data files were used to create a DIA chromatogram library (48) by searching against a Prosit-generated predicted spectral library (49) based on the UniProt_Mus musculus reviewed library_20191022. Mass spectrometry data for experimental samples were acquired in the Orbitrap using 8-m/z windows and searched against the chromatogram library. Mass spectrometry data was processed in Scaffold DIA (Proteome Software) filtered at 1% FDR using a minimum peptide count of 4. Normalized protein abundances were imported and wrangled using the *tidyverse* R package. Differential protein abundances were calculated with two-sample t-tests via the *rstatix* R package. KEGG enrichment analysis was performed using Enrichr. Data is shown as unadjusted p values for data exploration.

### Nanostring Gene Expression Analysis

RNA from lyophilized cerebrum and brainstem samples was extracted using the Maxwell^®^ RSC simplyRNA Tissue Kit/Instrument (Promega) with initial homogenization on a TissueLyser LT (Qiagen). RNA concentrations were determined using the Qubit^TM^ RNA Broad Range Assay Kit (Thermo Fisher Scientific). RNA Integrity (RIN), which ranges from 10 (highly intact) to 1 (strongly degraded), was assessed using a TapeStation 4150 via RNA ScreenTapes. 25 ng of RNA per sample (initial concentrations were 50-150 ng/μl with RIN 7-9), loaded into individual NanoString Hybridization reactions for 16 hours at 65°C, and loaded into a NanoString nCounter^®^ SPRINT cartridge/Analysis System. nSolver RCC files were imported into R using the read_rcc function from the *nanostringr* R package (50). Quality control was performed using the RUV_total function from the *NanoNormIter* R package to generate a continuous metadata value (“W1”) for each sample, which was used for correcting unwanted sources of sample-level variance (51). Differential gene expression analysis was performed using the *DESeq2* R package (52), controlling for sex and W1. Data was visualized using principal component analysis (PCA) for exploratory data analysis. Volcano plots were generated via the *EnhancedVolcano* R package using pairwise comparisons between genotypes and Bonferroni adjusted p-values. Heatmaps with hierarchical clustering and pathway analysis were generated via the *enrichR* R package (53) using unadjusted p values for data exploration.

### Immunohistochemistry

Sections were permeabilized in 0.5% Triton, blocked in 10% normal goat serum, and incubated with primary antibodies overnight at 4°C (pan-myeloid marker ionized calcium binding adaptor protein 1 (Iba1): WAKO, neuronal nuclei (NeuN): A60, EMD Millipore, myelin associated glycoprotein (MAG): D4G3, Cell Signaling, glial fibrillary acidic protein (GFAP): GA5, eBiosceince Invitrogen) and with secondary antibodies for 90 minutes (Invitrogen goat anti-rabbit AlexaFluor 555, goat anti-mouse AlexaFluor 488). Sections were mounted on permafrost slides, counterstained with DAPI, coverslips mounted and sealed. Slides were imaged using a Zeiss LSM 880 confocal microscope or a Bio-Tek Cytation 5 multi-mode imager. For image analysis, a region of interest (ROI) inclusive of the total brain section was selected and an ImageJ image processing analysis macro used to threshold the image and count particles and generate the area coverage and cell counts.

## Acknowledgments

This work was supported by the National Institutes of Health (NIH) [grant number AG066747 to SCH, AG013319-25S1 to JPP, GM130437 to JGR] and the Alzheimer’s Association [grant number AARG-21-846012 to SCH and JPP]. Lipidomics analyses were performed at the UT Health San Antonio Barshop Institute Functional Lipidomics Core partially supported by NIH [AG013319 (San Antonio Nathan Shock Center of Excellence in the Basic Biology of Aging), AG044271 (Claude D. Pepper Older Americans Independence Center), and AG061729 to XH]. Nanostring analysis was performed at the South Texas Alzheimer’s Center [AG066546].

## Notes

### Competing Interest Statement

The authors have declared no competing interest.

