## Supplemental Figures for "Multi-omics analyses reveal novel effects of PLCγ2 deficiency in the mouse brain"

**This PDF file includes:**

Supporting text

Figures S1 to S3

**Other supporting materials for this manuscript include the following:**

Datasets S1 to S8

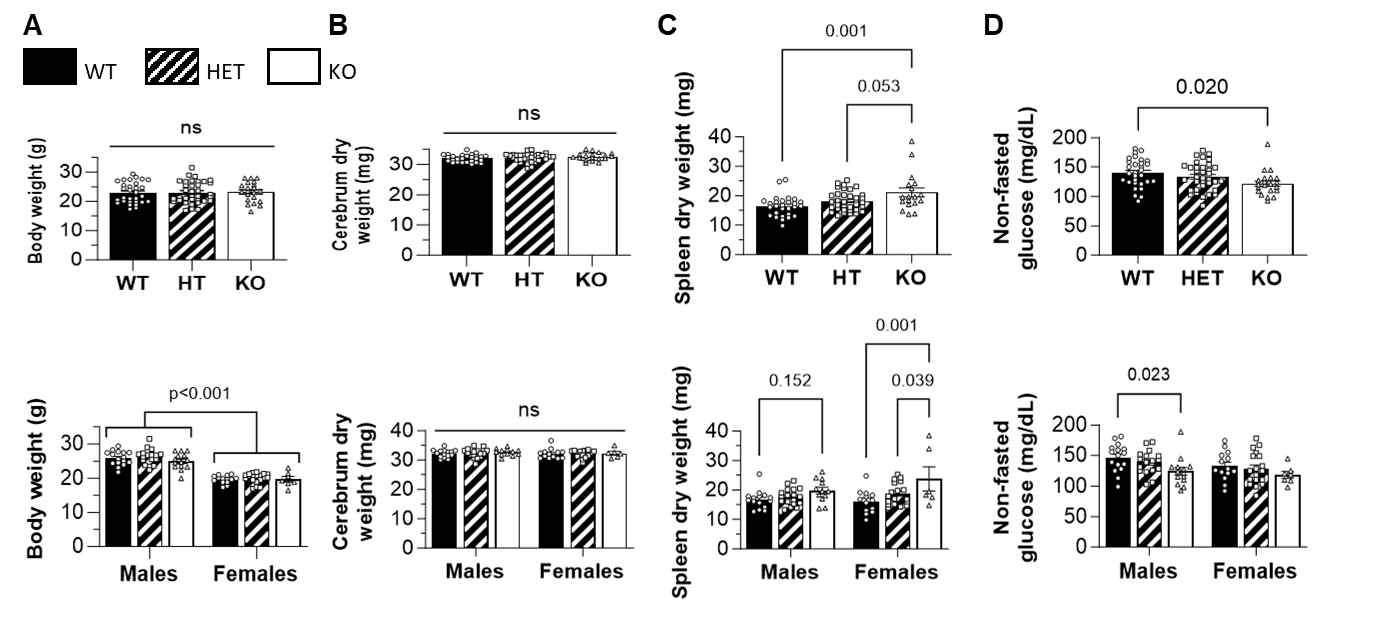

Fig. S1. Basic phenotyping of PLCG2 WT, HET, and KO mice with additional mice from another study.

To further explore the trending changes in non-fasting blood glucose found in Figure 1, we added data from additional control mice from another study; these data include both untreated mice and mice injected intraperitoneal with saline. A. Body weight. PLCG2 WT, HET, and KO mice show equivalent body weights with expected lower body weight in females. B. Brain weight. Brain dry weight was unchanged between genotypes or by sex. C. Non-fasting blood glucose. PLCG2 KO mice display a significant (18%) reduction in non-fasted blood glucose; post hoc analyses reveal significant reductions in male KO mice only. *p<0.05 *** p<0.001

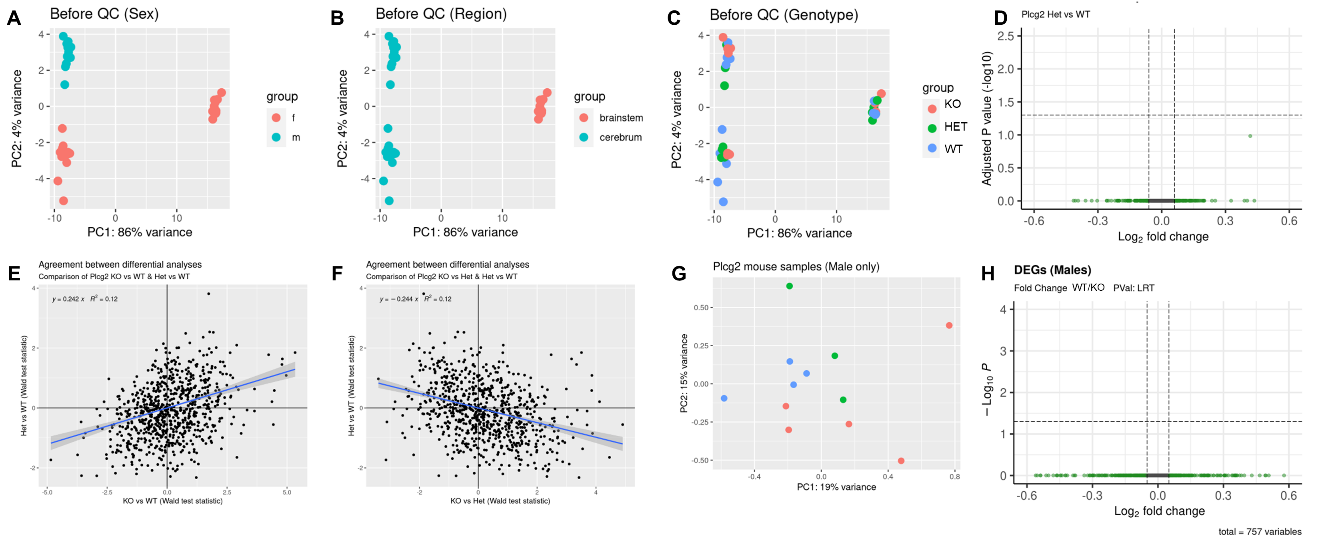

Fig. S2. Additional Nanostring gene expression analyses.

A. PCA plot shows sex-specific effects on gene expression before quality control/noise reduction. B. PCA plot shows brain region-specific effects on gene expression before quality control/noise reduction. C. Unbiased dimensionality reduction of the metadata did not reveal genotype effects. D. No DEGs were found between Plcg2 WT and HET mouse brains. E-F. Wald test statistics did not show correlation between E) KO vs. WT and WT vs. HET or F) KO vs. HET and HET vs. WT. G. Sex-specific PCA plots do not show clear separation between Plcg2 KO and Plcg2 WT/HET male brains. H. DESeq2 likelihood ratio test (LRT, Benjamini-Hochberg) shows no DEGs between WT and KO mice.

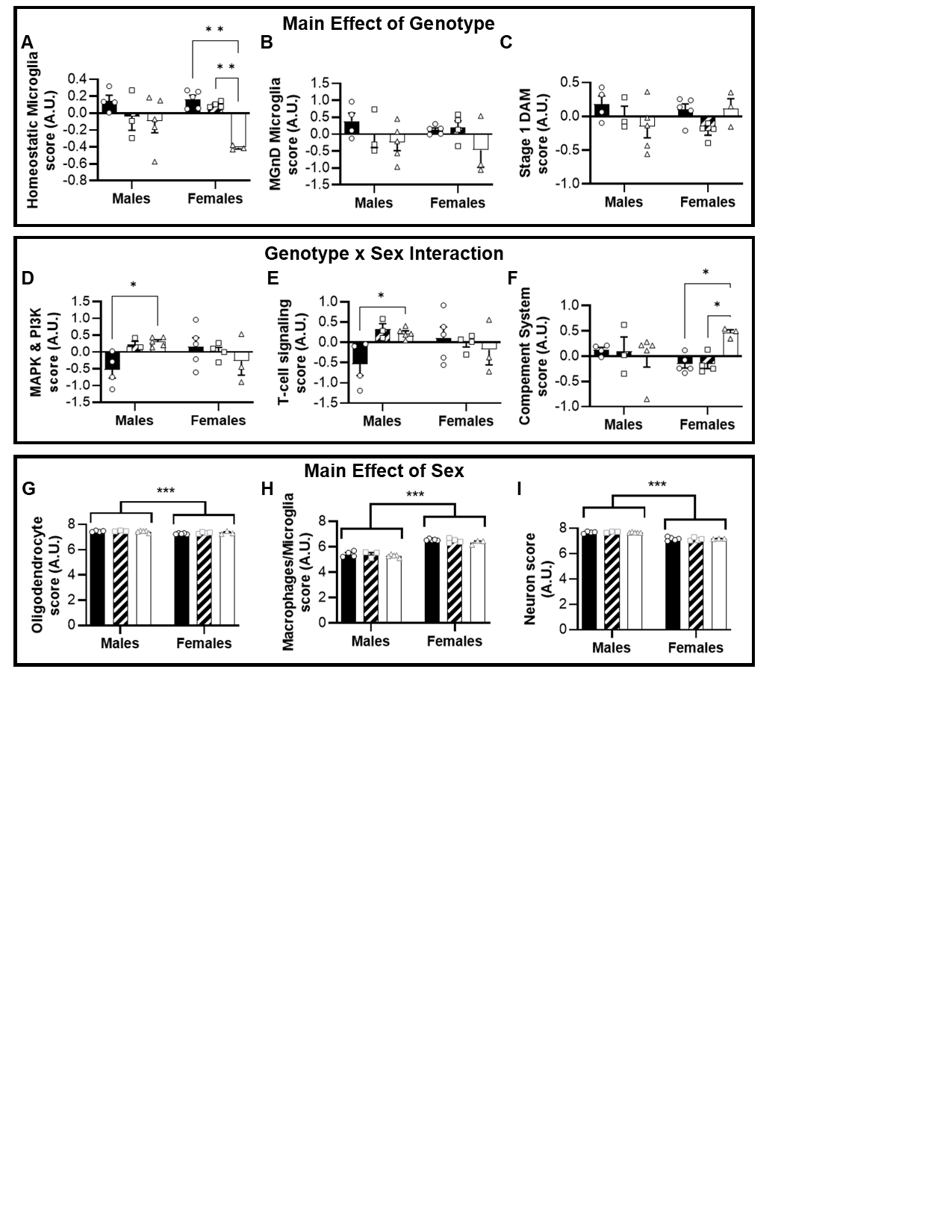

Fig S3. Nanostring Advanced Analysis

A-C. Difference in pathway scores showing significant main effects of genotype. A. Homeostatic microglia pathway scores are significantly reduced in female PLCG2 KO mice. B. MGnD pathway scores. C. DAM stage 1 pathway scores. D-F. Difference in pathway scores showing significant genotype by sex interactions. D. MAPK and PI3K pathway score is significantly different between male WT and KO mice. E. T cell signaling pathway score is significant different between male WT and KO mice. F. Complement pathway score is significantly higher in PLCG2 KO female mice compared to WT and HET mice. G-I. Difference in pathway scores showing a significant main effect of sex. G. Oligodendrocyte genes are slightly reduced in females. H. Macrophage/microglia genes are reduced in males. I. Neuron genes are slightly reduced in females.

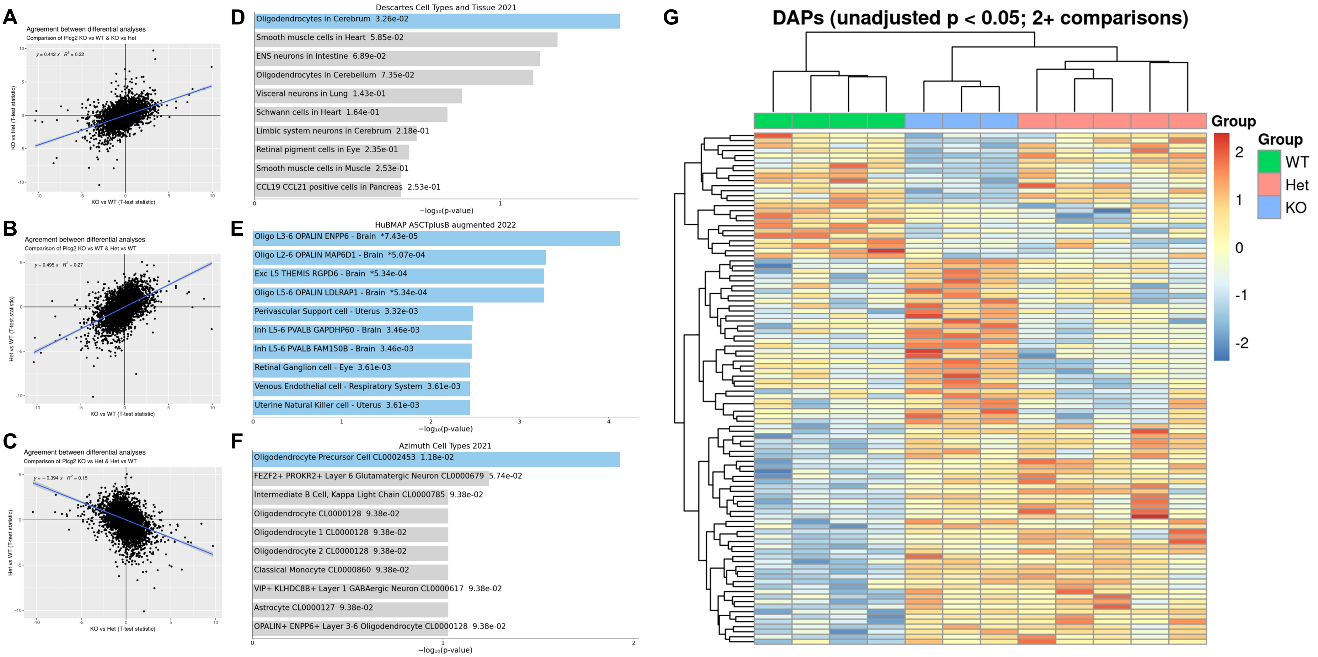

Fig S4. Proteomics

A-C. Wald test statistics showed KO vs WT and HET vs WT were similar, suggesting that the KO and Het genotype conditions are similar to one another at the protein level. A) KO vs. WT and KO vs. HET B) KO vs. WT and HET vs. WT C) KO vs HET and HET vs. WT. D-F. Enrichr Cell type analyses using underabundant proteins revealed oligodendrocytes and oligodendrocyte precursor cells are significantly altered (q-value<0.05) in KO brains. G. Differentially abundant proteins (DAPs) between WT, HET, and KO mice. All DAPs with under an unadjusted p < 0.05 in at least two comparisons are shown.

Dataset S1 (separate file). Lipidomics analysis – males

Dataset S2 (separate file). Lipidomics analysis – females

Dataset S3 (separate file). Nanostring Glia Profiling Panel DEGs for PLCG KO vs. WT

Dataset S4 (separate file). Nanostring Glia Profiling Panel DEGs for PLCG KO vs. HET

Dataset S5 (separate file). Nanostring Glia Profiling Panel DEGs for PLCG HET vs. WT

Dataset S6 (separate file). Nanostring Glia Profiling Panel DEG LRT analysis by sex

Dataset S7 (separate file). Proteomics DAPs for PLCG KO vs. WT

Dataset S8 (separate file). Proteomics DAPs for PLCG HET vs. WT

Dataset S9 (separate file). Proteomics DAPs for PLCG KO vs. HET
